## Supplementary Information for "Statistical Mechanical theory for spatio-temporal evolution of Intra-tumor heterogeneity in cancers: Analysis of Multiregion sequencing data"

(Dated: February 20, 2022)

*

### I. TUMOR MUTATION BURDEN (TMB) FROM M-SEQ DATA

Multi-region sequencing (M-seq), which has provided the most direct evidence for pervasive signature of Intratumor heterogeneity (ITH) in solid tumors, is used to delineate the distribution of mutations in different regions of a single solid tumor [1–6]. Figure S1 shows a schematic representation of a typical M-seq data. The data is in the form of a matrix where the rows are the names of the genes and the columns label the biopsied tumor regions. In Figure S1, the mutated genes are in yellow, and blue denotes the non-mutated genes. Usually the mutations, sequenced using M-seq, are non-silent or non-synonymous [2–4, 6], which implies that they impact the tumor phenotype. From the M-seq data, we can classify mutations as public, branched or private. Public genes are mutated in all the sequenced regions. If a mutation is present in more than one region but not in all, then it is referred to as branched. If a mutated gene is found in only one region, then it is classified as private [7]. The presence of a large number of branched and private mutations, enhances ITH.

We first analyzed the M-seq data for four cancer types from different patients in order to assess the extent of ITH. Three of them (skin, lung and esophagus) are predominantly caused by exogenous factors while the fourth ( kidney) is linked to endogenous factors [8]. The data for skin cancer was taken from Harbst et al.[3], lung from Debruin et al.[4], esophagus from Cao et al.[6]. For kidney cancer, we used data from the experiments by Gerlinger et al.[2]. The number of patients, whose tumors were sequenced, are eight[3], seven[4], two[6] and ten[2] for skin, lung, esophagus and kidney cancer respectively. In some sense, the sample sizes are small but are sufficient to calculate ITH using our theory, and establish its efficacy.

Figure S2 shows the number of mutations, referred to as tumor muation burden (TMB), in different regions of tumors across eight skin cancer patients. The average number of mutations among the eight patients is $\approx 464$. The maximum number of mutations was found in patient Mm1749 who acquired $\approx 779$ mutations per tumor region. In contrast, patient Mm1837 has the smallest number (36 per tumor region) of mutations. Figure S3 summarizes the results for the six lung cancer patients. The average number of mutations among the six lung cancer patients is $\approx 238$. The maximum (minimum) number of mutations was found in patient L011 (patient L008) with $\approx 375$ ($\approx 98$) mutations per tumor region. Figure S4 shows the number of mutations in two patients (A and B) with esophageal cancer. Patient A had $\approx 71$ mutations per tumor region whereas patient B had $\approx 91$ mutations per tumor region. Figure S5 shows the number of mutations in ten kidney cancer patients. The average number of mutations across these ten patients was $\approx 43$. Patient EV001 with $\approx 67$ mutations per tumor region had the maximum whereas patient RK26 with $\approx 21$ mutations per region had the minimum number of mutations.

The data shows that cancers caused predominantly due to exogenous mutagens have more mutations. These results are consistent with previous studies [9, 10]. However, strong claims about esophageal cancer should be made with caution because M-seq data is available only for two patients.

### II. RELATIONSHIP BETWEEN ITH AND TMB

The tumor mutation burden, TMB, is defined as the number of mutations accumulated over a time period, $t$. In the M-Seq data, the average number of mutations per region is $(1 - x_t)n$, where $n$ is the length of the string. Because $x_t = \frac{1 \pm \sqrt{1-4H'}}{2}$, one can derive a relationship between TMB and ITH using $H' = \frac{ITH}{2F(\alpha)}$. The relationship is given by $TMB = n[\frac{1 \mp \sqrt{1-4H'}}{2}]$. One can also estimate the number of single nucleotides mutated per year from the M-Seq data ($M_{SN}$) by using the relationship,

$$G(1-(1-p_D)^t) = n(1-x_t), \tag{S.1}$$

where $p_D$ is the mutation probability of a gene per year, G is total number of genes in the genome ($\approx 20,000$), and $t$ is the time from tumor initiation to detection. For skin and kidney cancers, $t$ is about 13, and 20 years, respectively[11]. For lung cancer, the TMB is found to show a weak correlation with the patient age, therefore, the mutation rate is not detected in Ref. [11]. Here, we estimated the $M_{SN}$ for skin and kindey cancer patients. Using, $M_{SN} = (\frac{p_D}{N_G})(N_D) = \frac{(p_D)}{(10^4)}(3 \times 10^9)$, where $N_G$ ($\approx 10^4$ $bps$) is the average number of nucleotides in a gene (10,000) and $N_D$ is the number of nucleotides on the DNA. Figure S10 shows the plot for $M_{SN}$ for the two cancer types: skin, and kidney. For skin and kidney cancers, the mean $M_{SN}$ is $\sim 1200$ and $\sim 125$ respectively, which is consistent with our previous findings, Skin ($\sim 1500$) and Kidney ($\sim 70$), from The Cancer Genome Atlas (TCGA) data (see Figures 1 (Kidney) and 3 (Skin) in [11]).

### III. TUMOR SIMULATION MODEL

We created a three-dimensional (3D) cellular automaton (CA) model, similar to a computational model studied previously in a pioneering paper [12], to calculate ITH in order to not only validate the theory but provide mechanistic insights into the spatio-temporal propagation of ITH. The 3D space is divided into finite lattice sites of size $N \times N \times N$, with $N = 100$. A lattice site, whose position is represented as $[i, j, k]$, is either vacant ($L[i, j, k] = 0$) or occupied ($L[i, j, k] = 1$). A lattice site can only be occupied by a one cell (see Fig S11a).

In the process of evolution, a normal cell may become cancerous by accumulating non-synonymous (or non-silent) mutations. The number of non-synonymous mutations, $n_d[i, j, k]$, at position $[i, j, k]$, characterizes whether a cell is normal or cancerous. If $n_d[i, j, k] = 0$, then the cell is normal otherwise it is cancerous (see Fig. S11b).

We model the DNA representing the mutation sites to calculate the ITH. Each cell contains DNA in the form of an information string, $X$, of length $n$. A site in $X$ is 0 (no mutation) or 1 (mutated). For simplicity, we consider the same fitness advantage, $s_d$, for all the non-synonymous mutations. We assume $s_d = 0$, unless stated otherwise. The assumption that $s_d = 0$ implies that evolution is neutral. As a result, there is no fitness advantage upon mutation.

Cell division is controlled by birth probability ($\alpha$) provided neighboring lattice site (see Fig. S11c) is vacant. In 3D, the neighbors are defined by the 26 lattice sites around a cell. The birth probability of a cell, $\alpha[i, j, k]$, is given by the relation,

$$\alpha[i, j, k] = \alpha_o[i, j, k] + s_d \times n_d[i, j, k], \tag{S.2}$$

where, $\alpha_o[i, j, k]$ is the initial birth probability of the cell. When a cell divides, we copy the mutation profile of the parent cell ($P$) to its daughter cell (D) ( $X^P = X^D$). At each time step (equivalent to one cell generation), a cell may acquire non-synonymous mutations with probability $p$ per site. We also assume that there is no back mutation. A cell can also undergo apoptosis with probability $\beta[i,j,k] = 1 - \alpha[i,j,k]$ (see Fig. S11d).

**Evolution with selection:** To assess the robustness of the conclusion that ITH is pervasive and varies greatly within a single tumor, we also carried out simulations with $s_d \neq 0$. The effect of fitness advantage is modeled by increasing the birth rate given in S.2. Figure S12, shows that $s_d$ plays a role in the range, $0.3 \leq x_t \leq 0.7$. For $s_d = 0.0001$ and 0.001, we see that the ITH remains approximately the same as in the case of $s_d = 0$. However, for the case of $s_d = 0.01$, we see that heterogeneity increases slightly in the domain $0.3 \leq x_t \leq 0.7$. This occurs because as we increase $s_d$, the birth probability increases as a function of time, and hence it will result enhance heterogeneity. We know that at higher birth probability, there is more heterogeneity as explained in the main text.

**Initial Conditions:** To initiate the simulations, we introduce a single normal cell at the center of the 3D lattice. This cell has a birth probability $\alpha[\frac{N}{2}, \frac{N}{2}, \frac{N}{2}] = \alpha_o$ without any non-synonymous mutations ($n_d[\frac{N}{2}, \frac{N}{2}, \frac{N}{2}] = 0$). Following the evolutionary rules described above, this cell divides and acquires mutations stochastically resulting in a tumor mass with heterogeneous DNA strings.

**Termination of Simulations:** We stop the simulations once the size of the tumor reaches $M$ cells. We assume that $M = 50,000$, if not mentioned explicitly. The simulations were performed using an in house code using the MATLAB software.

TABLE I: Hamming distance (HD) between the pairs of columns for the data in Figure 1b of the main text.

| Pairs | Hamming Distance(HD) |
|---|---|
| (R1, R2) | 0.13 |
| (R1, R3) | 0.10 |
| (R1, R4) | 0.07 |
| (R1, R5) | 0.04 |
| (R1, R6) | 0.31 |
| (R1, R7) | 0.18 |
| (R2, R3) | 0.08 |
| (R2, R4) | 0.06 |
| (R2, R5) | 0.12 |
| (R2, R6) | 0.30 |
| (R2, R7) | 0.22 |
| (R3,R4) | 0.02 |
| (R3, R5) | 0.08 |
| (R3, R6) | 0.27 |
| (R3, R7) | 0.18 |
| (R4, R5) | 0.06 |
| (R4, R6) | 0.24 |
| (R4, R7) | 0.16 |
| (R5, R6) | 0.30 |
| (R5, R7) | 0.17 |
| (R6, R7) | 0.40 |

.

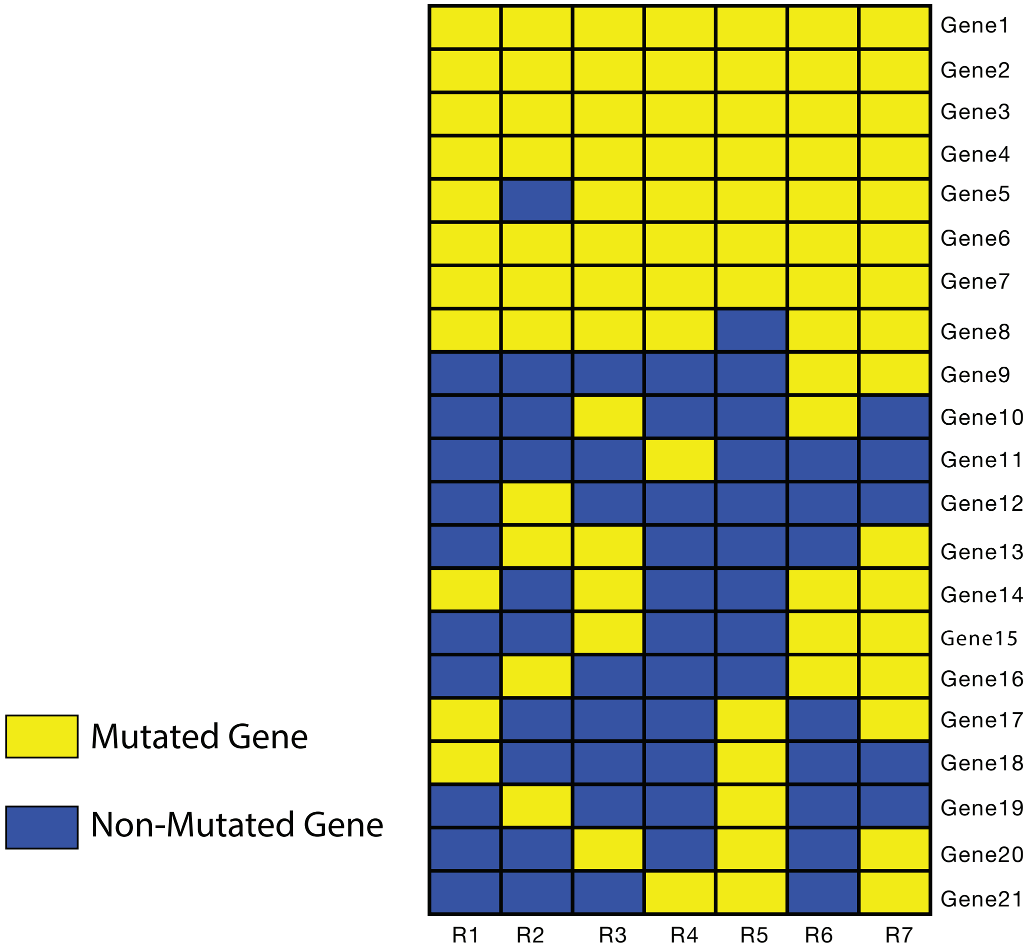


**Figure** S1: Schematic matrix representation of a typical mutiregion sequencing (M-seq) data depicting region-specific distribution of mutations within a single tumor. The rows denote the gene names ( Gene1 to Gene21), and the columns are the regions of the sequenced tumor ( R1 to R7). The yellow (blue) boxes represent a mutated (non-mutated) genes. In the experiments, the number of regions sequenced may vary depending on the size and quality of the tumor sample. The number of genes mutated also changes across various cancer types. Note that mutations in Gene 1, Gene 2, Gene 3 are public, the mutations in Gene 11 is private, and an example of branched mutation is Gene 20.

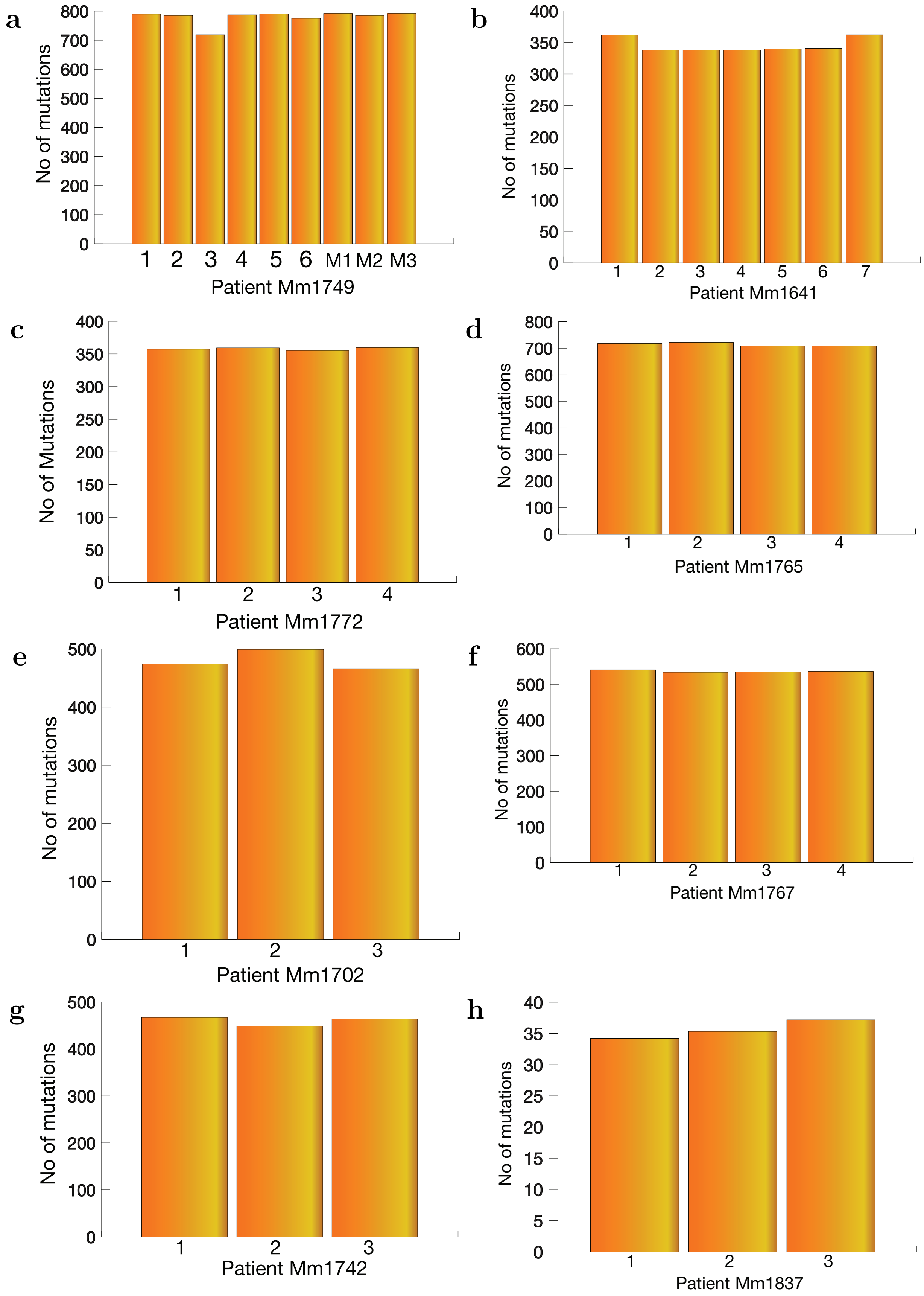


**Figure** S2: Number of mutations in different regions of tumors for eight skin cancer patients. The number of mutations varies from patient to patient.

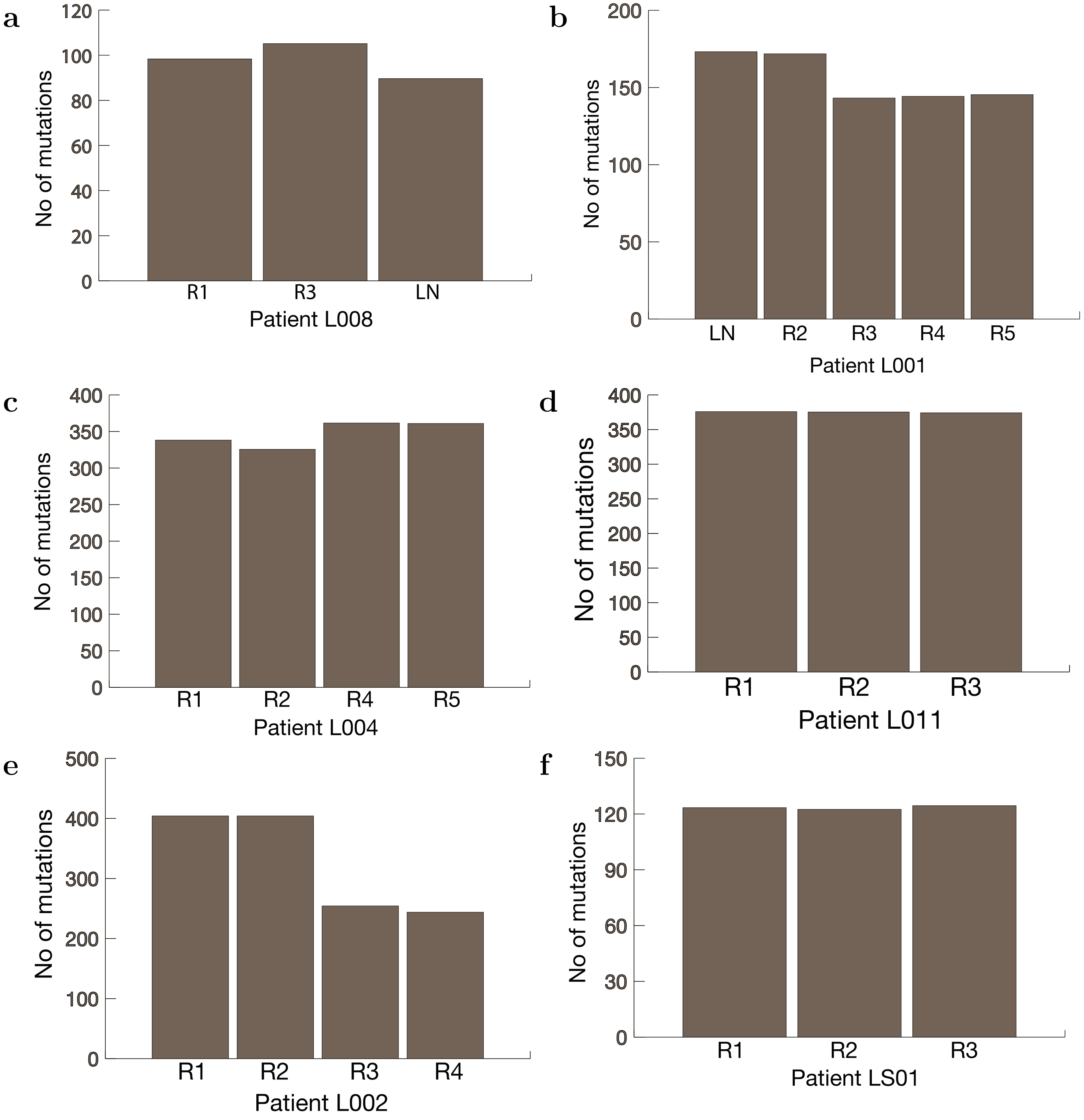


**Figure** S3: Number of mutations in different regions of tumors for six patients with lung cancer. The average number of mutations among the six lung cancer patients was found to be $\approx 238$. The maximum number of mutations was found in patient L011 with $\approx 375$ mutations per tumor region. The minimum number of mutations was found in L008 with $\approx 98$ mutations per tumor region.

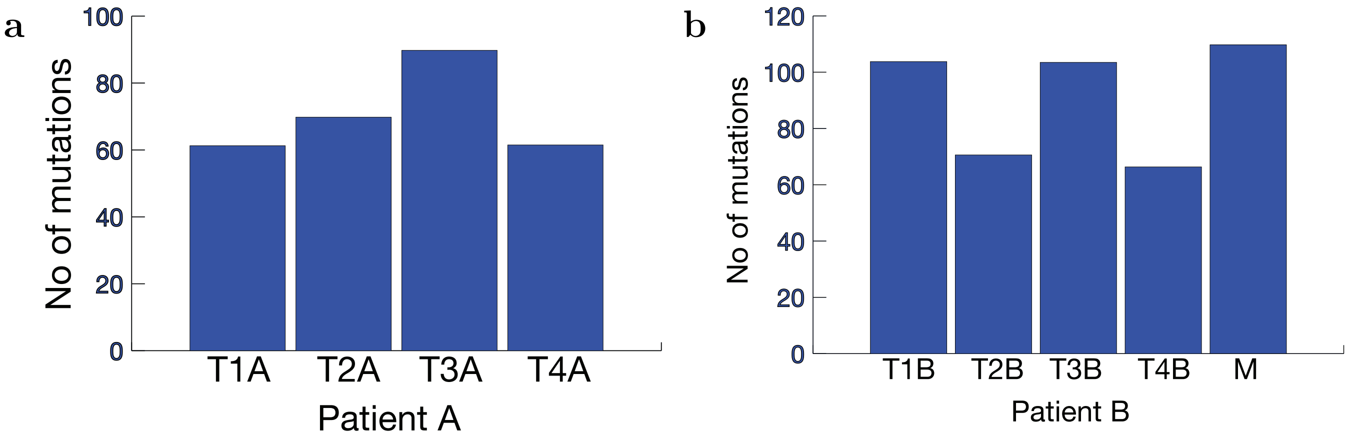


**Figure** S4: Number of mutations in different regions of tumors for two esophageal cancer patients (A and B). Patient A had $\approx 71$ mutations per tumor region whereas patient B had $\approx 91$ mutations per tumor region.

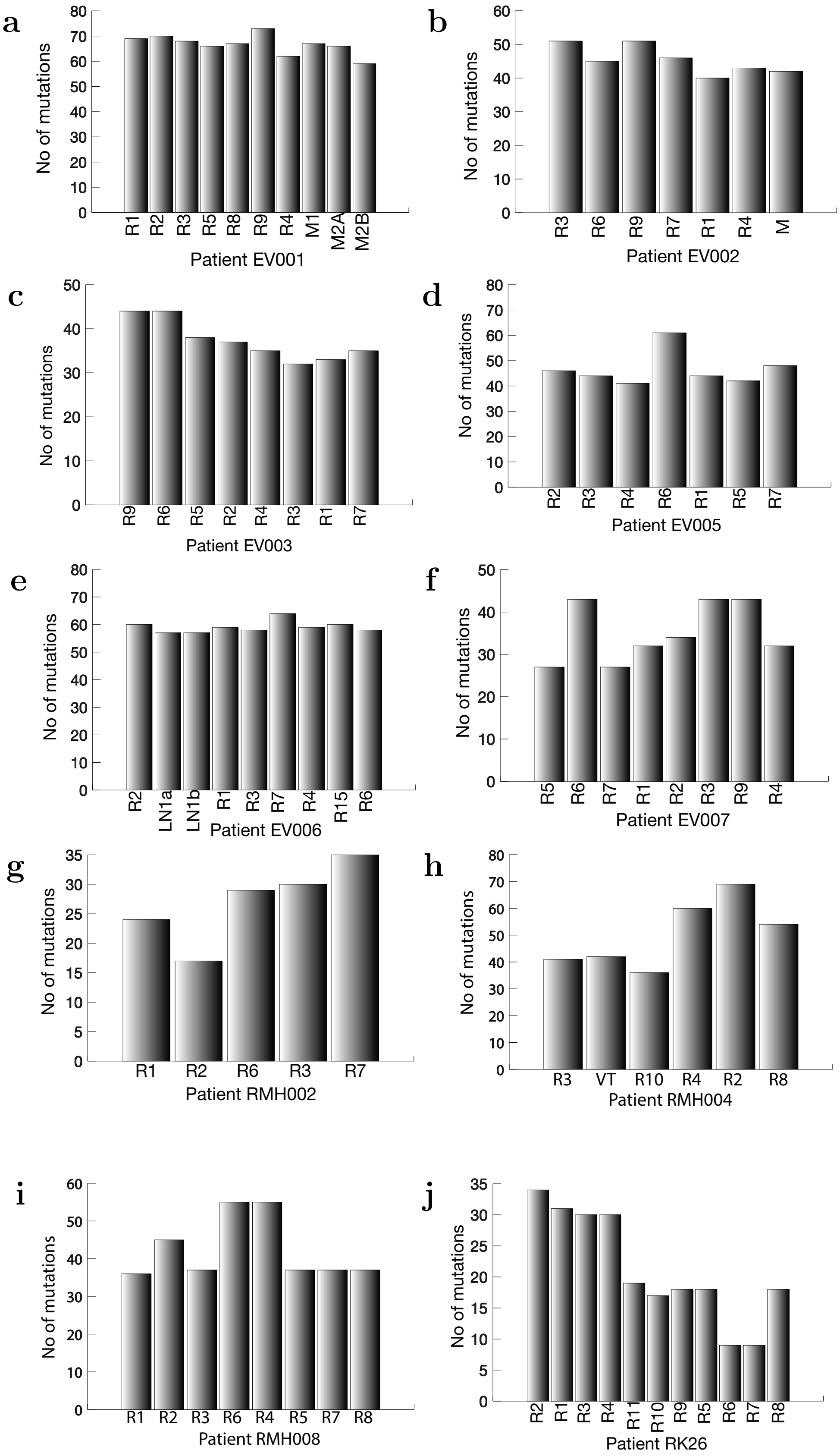


**Figure** S5: Number of mutations in different regions of tumors across ten patients with kidney cancer. The average number of mutations across these ten patients was $\approx 43$. Patient EV001 with $\approx 67$ mutations per tumor region had the maximum whereas patient RK26 with $\approx 21$ mutations per region had the minimum number of mutations.

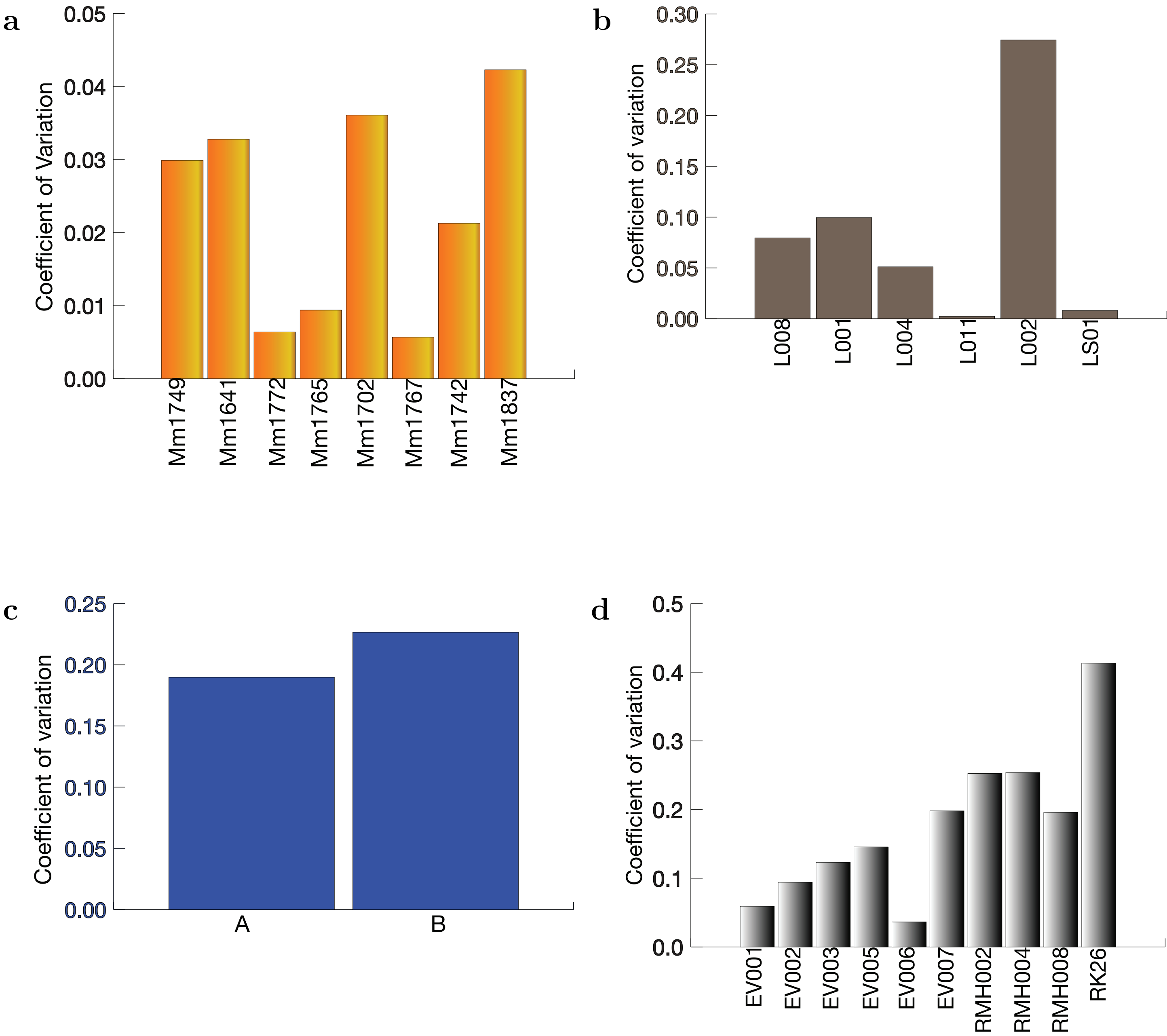


**Figure** S6: Coefficient of variation ($c_v$), defined in Eq.(10) in the main text, for exogenous (skin, lung and esophagus) and endogenous (kidney) cancers. **(a)** $c_v$ for eight skin cancer patients. The $c_v$ varies from 0.042 (patient Mm1837) to 0.006 (patient Mm1767). **(b)** Same as (a) except for six lung cancer patients. $c_v$ varies from $\approx 0.002$ (patient L011) to $\approx 0.27$ (patient L002). **(c)** shows the $c_v$ for the two esophageal cancer patients (A and B). **(d)** Same as (a) except for the ten kidney cancer patients. In this case, $c_v$ changes from $\approx 0.004$ (patient EV006) to 0.4 (patient RK26).

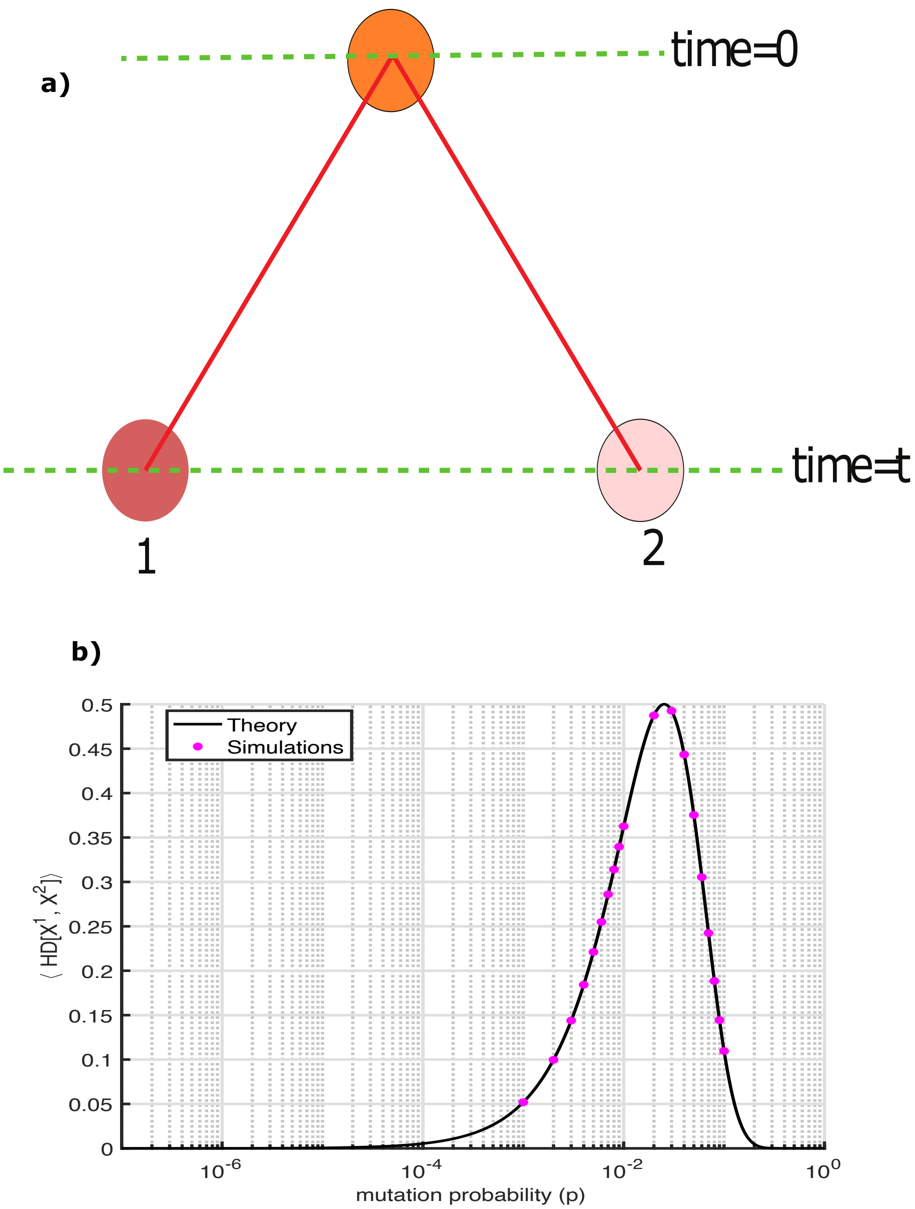


**Figure** S7: **Independent Evolution: (a)** Evolution of two cells (brown and pink). The two cells are normal at time $t = 0$ (orange), and evolve independently acquiring exogenous mutations. **(b)** The average heterogeneity between the two cells ($\langle HD[X^1, X^2] \rangle$) that evolve independently as a function of the mutation probability (p). The dots are simulation results carried out for 27 time steps or generations. The simulation was averaged over $\approx 4 \times 10^8$ pairs of cells. The black line is calculated using the theoretical prediction given in Eq. (18) in the main text.

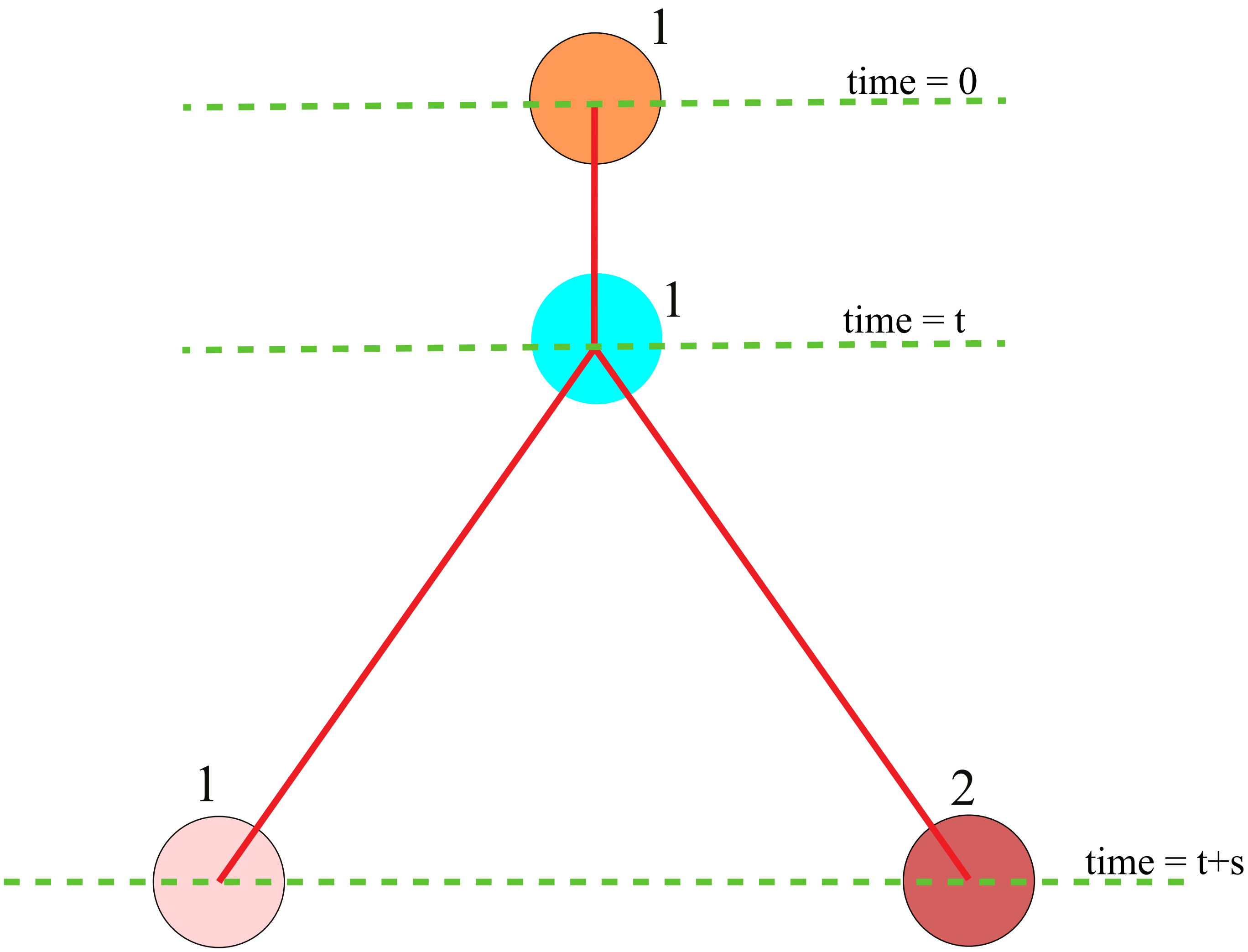


**Figure** S8: **Schematic of correlated (dependent) evolution of 2 cells (black filled circles)**. At time $t = 0$, evolution is initiated with a single normal cell (cell 1). It evolved till time $t$, gathering exogenous mutations. At time $t$, cell 1 gives birth to cell 2. After time $t$, the 2 cells evolve independently for the next $s$ time steps. The ancestor of cell 2, at $t + s$, is cell 1. Therefore, we consider them to have undergone correlated evolution. At $t$, the genetic information in the blue cell is copied to the pink and brown cells. Because the pink cell at $t = t + s$ is genotypically identical to the blue cell, we use the same label.

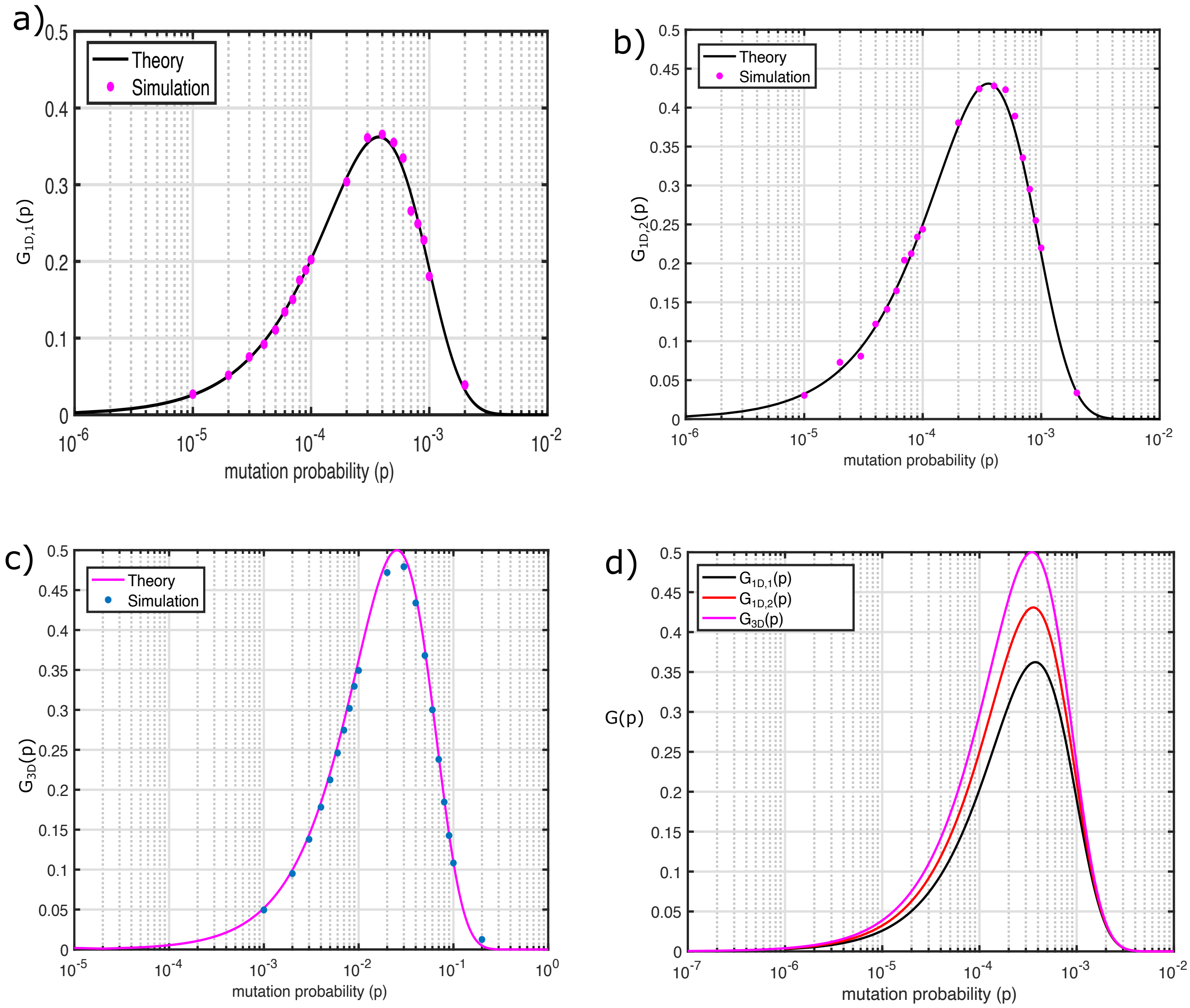


**Figure** S9: (Continued on the next page)

**Figure** S9: **Comparison between theory and simulations for G(p) as a function of $p$ for different geometries.** In all cases $\alpha = 1$, dots are results from simulations, and lines are theoretical predictions. Eqns. (25), (26) and (28) in the main text, plotted on the y-axis are evaluated at $\tau_1 = 2,000$ steps (generations) in (a) and (b) and $\tau_2 = 27$ generations in (c)). For clarity we do not show the time argument in G(p, t). **(a)**Average heterogeneity within an evolving tumor in the 1D semi-infinite lattice, $G_{1D,1}(p)$, for $\alpha = 1$. The dots are generated using simulations with $\alpha = 1$. The number of cells at the end of simulations is $2,000$. The black line corresponds to equation (25) in the main text. Due to the correlated nature of evolution, the heterogeneity is smaller compared to the independent evolution of cells ( Figure S7b). **(b)** Average heterogeneity within an evolving tumor in 1D infinite lattice, $G_{2D,1}(p)$. The number of cells at the end of the imulation is $4,000$. The black line is calculated using Eq. (26) in the main text. Due to temporal correlations present among the cells, the heterogeneity in this case is smaller compared to the case of independent evolution of cells (i.e Figure S7b). **(c)** Same as (b) except the results for a tumor in 3D lattice, $G_{3D}(p)$. The number of cells at the end of the simulation was $\approx 50,000$. Magenta line is a plot of Eq. (28) in the main text. The behavior is similar to cells evolving independently because in 3D, the number of branches in an evolutionary tree is very large ($\propto A_c$) where $A_c$ surface area of the tumor. **(d)** Comparison of average heterogeneity, $G(p)$, in 1D semi-infinite (black), 1D infinite (red) and 3D lattice (magenta). $G_{1D,1}(p)$ has the smallest value, $G_{1D,2}(p)$ has an intermediate value and $G_{3D}$ for the 3D lattice has the maximum value. The plots illustrate that heterogeneity increases as the degree of branching within an evolutionary tree increases.

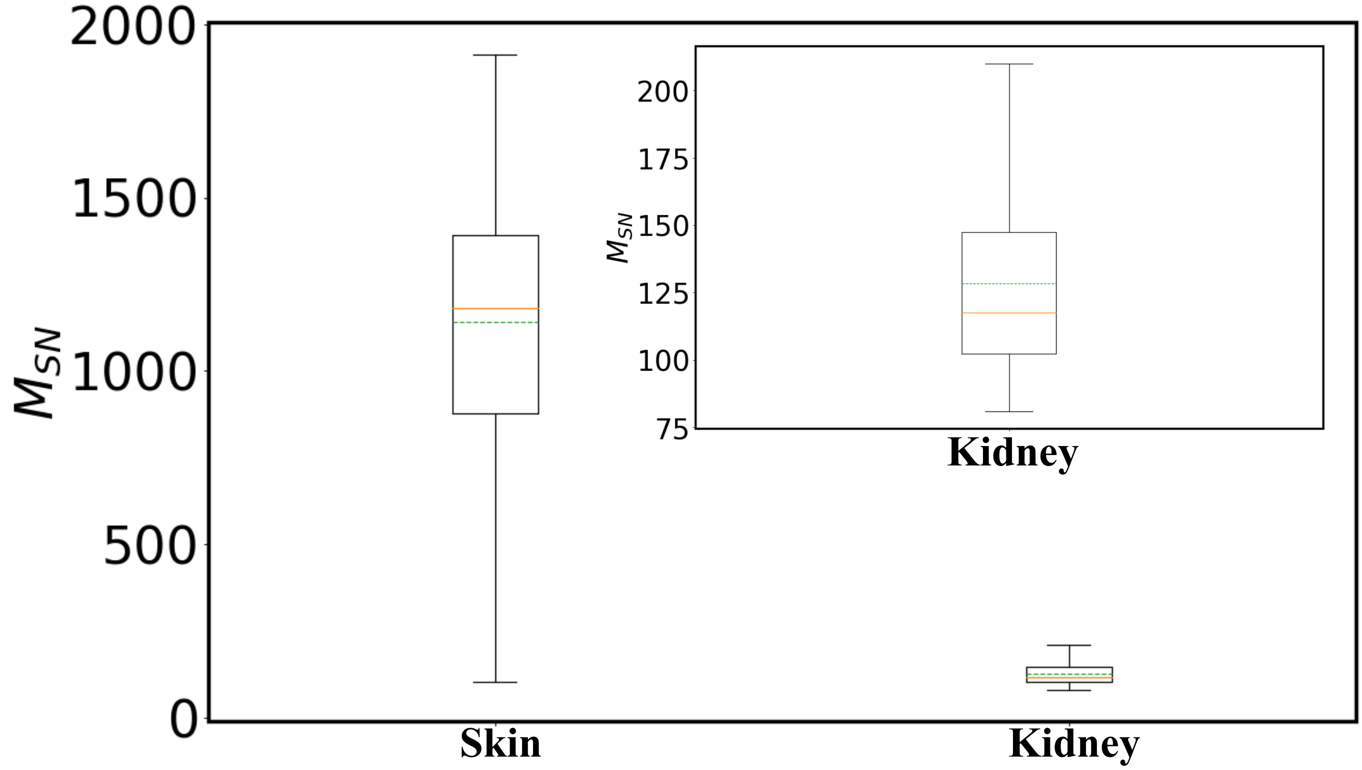


**Figure** S10: Number of single nucleotide mutations per year ($M_{SN}$) for the skin, kidney cancer. The yellow (green) line is the median(mean) of the data. The inset shows the box plot for kidney cancer.

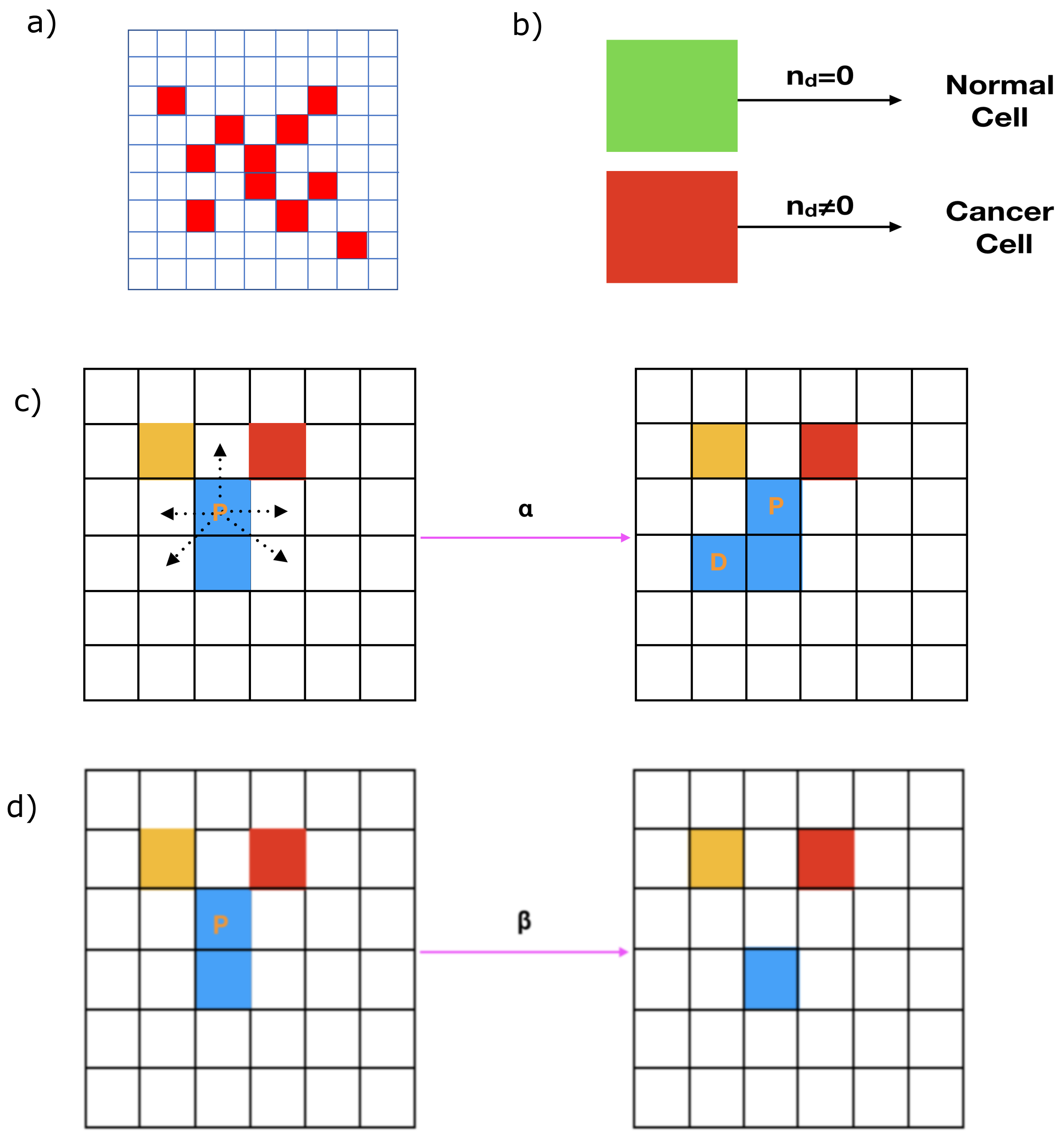


**Figure** S11: **Schematic of the simulation framework**. **(a)** Cross-section of a 3D ($N^3$) lattice model for cancer evolution. For purposes of illustration, we choose $N = 9$. The red colored squares denote an occupied site ($L[i, j, k] = 1$) and white squares denote the site is vacant ($L[i, j, k] = 0$).**(b)** Cartoon depicting a cancer cell and a normal cell. A cell with zero number of non-synonymous mutations ($n_d = 0$) is normal. It is cancerous otherwise ($n_d > 0$). (c),(d) Cartoon depicting cell division and apoptosis. **(c)** Shows that the parent cell (P) divides (with probability $\alpha[i, j, k]$), and a daughter cell (D) is born. The D cell occupies one of the available sites (indicated by the five dashed arrows). In **(d)** the parent cell (P) undergoes apoptosis (with a probability $\beta[i, j, k] = 1 - \alpha(i, j, k)$), and the site becomes vacant. In both the figures, different colors means that cells have different genetic composition.

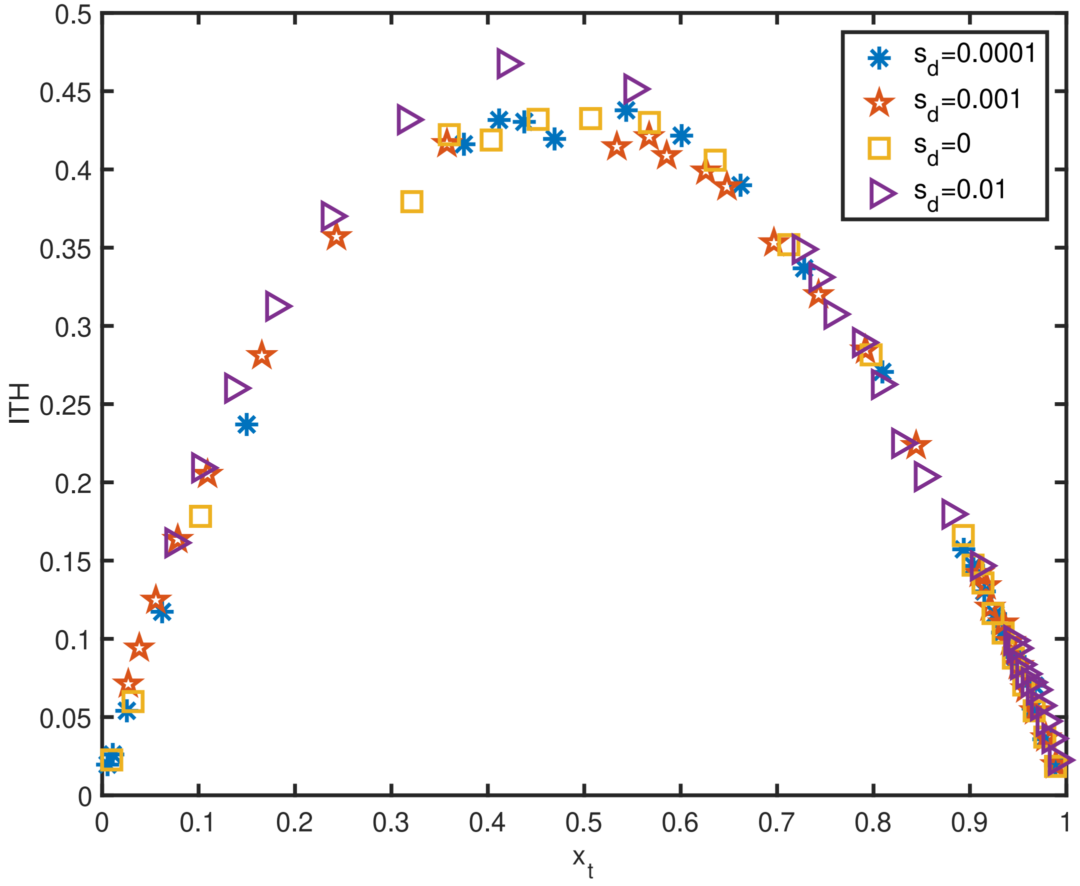


**Figure** S12: **Role of fitness advantage ($s_d$) on ITH**. ITH as a function of $x_t$ for three non-zero $s_d$ values. For comparison, we also plot the results for $s_d = 0$. For $s_d = 0.0001$ and $s_d = 0.001$, the ITH is approximately similar to case of $s_d = 0$. However, for $s_d = 0.01$, we see noticeable deviations from the master curve in the range $0.3 \leq x_t \leq 0.7$. The results were obtained for $\alpha = 0.55$ and $t$ when the number of cells is $50,000$.

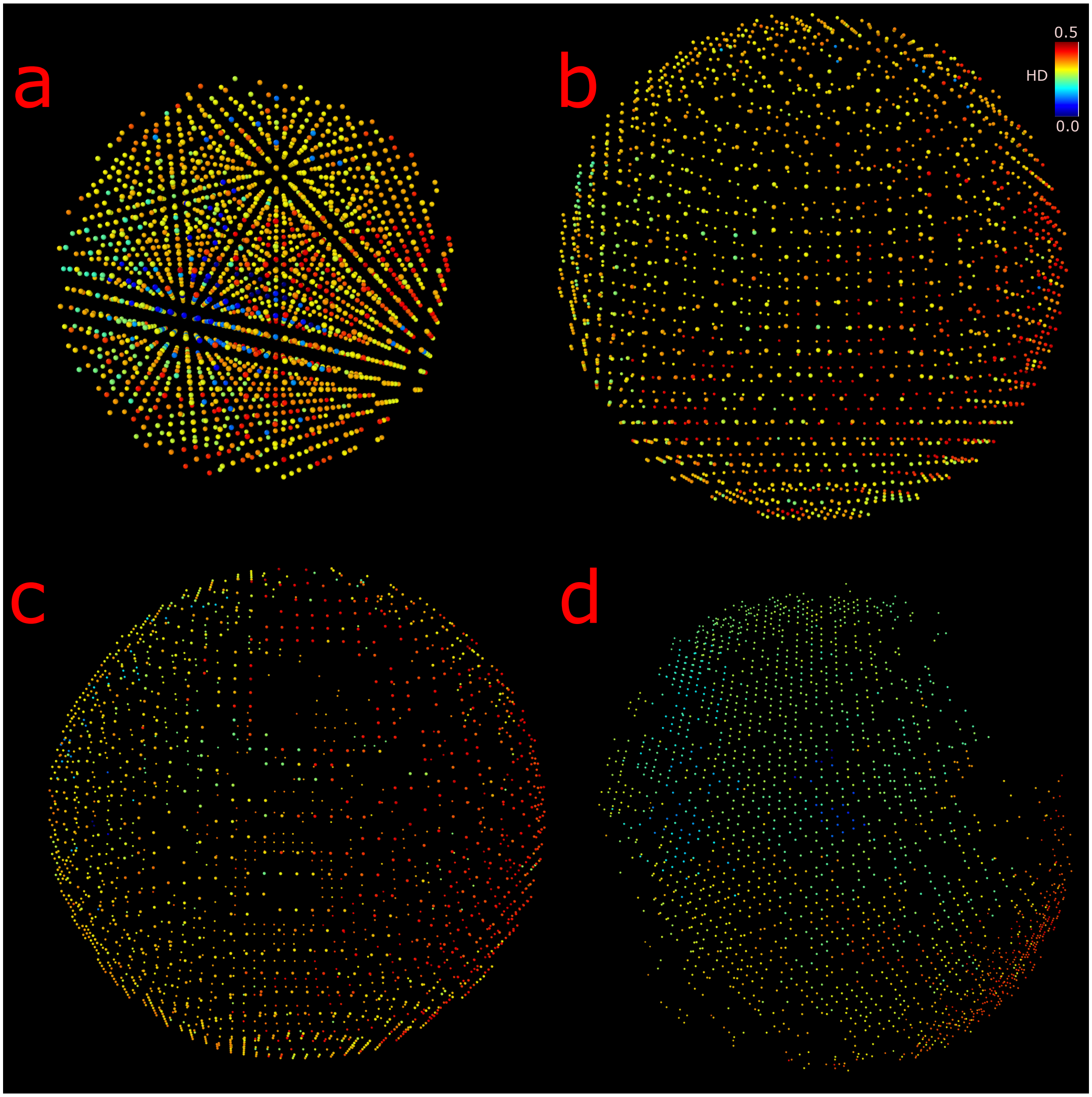


**Figure** S13: **Sub-sample to sub-sample variation in ITH in different regions of a simulated tumor**. The images were generated using 3D lattice simulations with $\alpha = 0.55$ and $p = 0.003$. The image corresponds to tumor evolution that was terminated after $t = 113$ generations. In all the cases, 2, 000 cells are present and the HD value is calculated with respect to the cell in the center. **(a)** HD values for 2, 000 cells located closest to the center, depicted by their color. **(b)** Same as (a) except for cells located approximately 15 lattice units from the center (same as Figure 8 in main text). **(c)** Same as (a) except for cells located approximately 20 lattice units from the center. **(d)** Same as (a) except for cells located approximately 27 lattice units from the center. The scale on the top right corner gives the scale for HD.

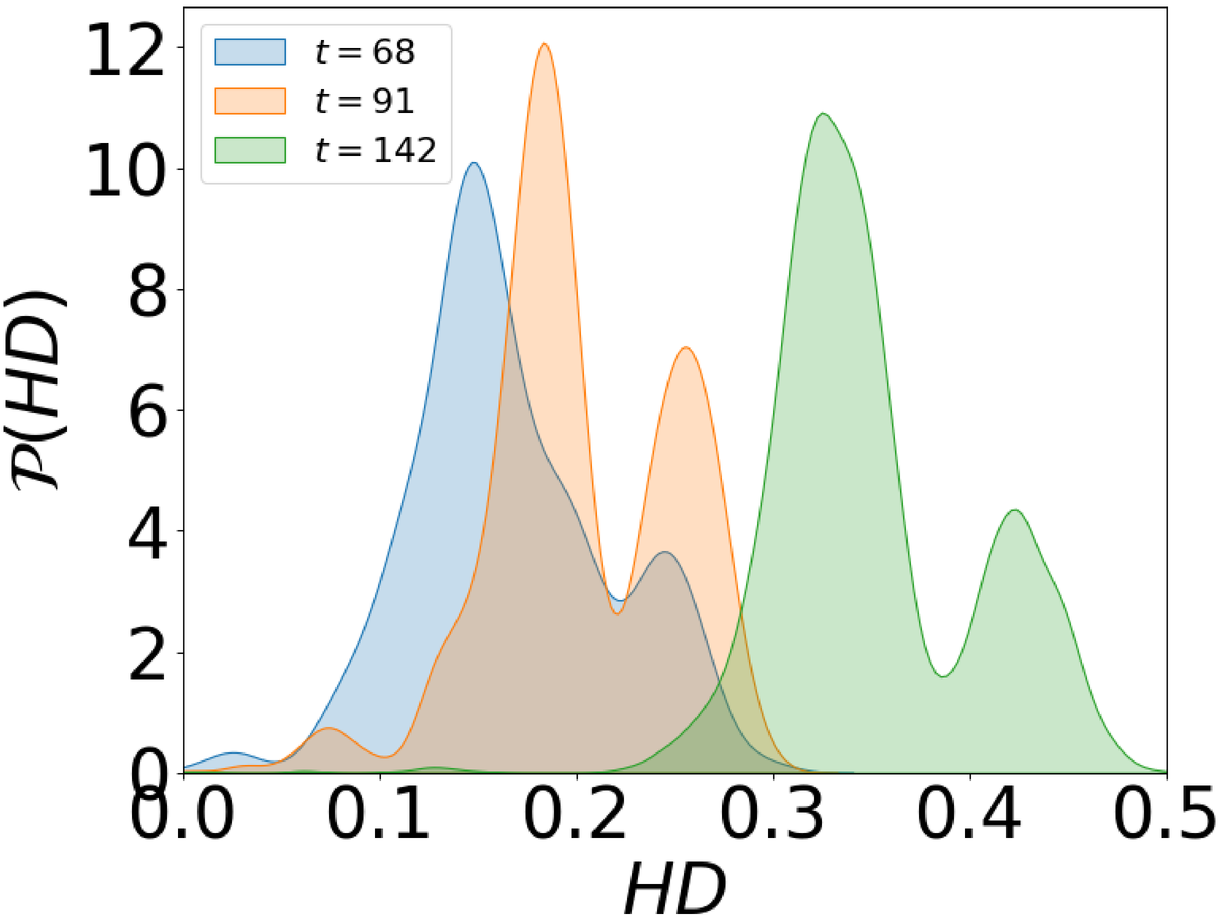


**Figure** S14: **Time evolution of the Hamming distance probability distribution** $\mathcal{P}(HD)$. From left to right, $\mathcal{P}(HD)$ corresponds to $t = 68, 91$ and 142 generations. All the distribution were generated using 3D lattice simulations with $\alpha = 0.55$ and $p = 0.003$. HD values were calculated with respect to the center cell.

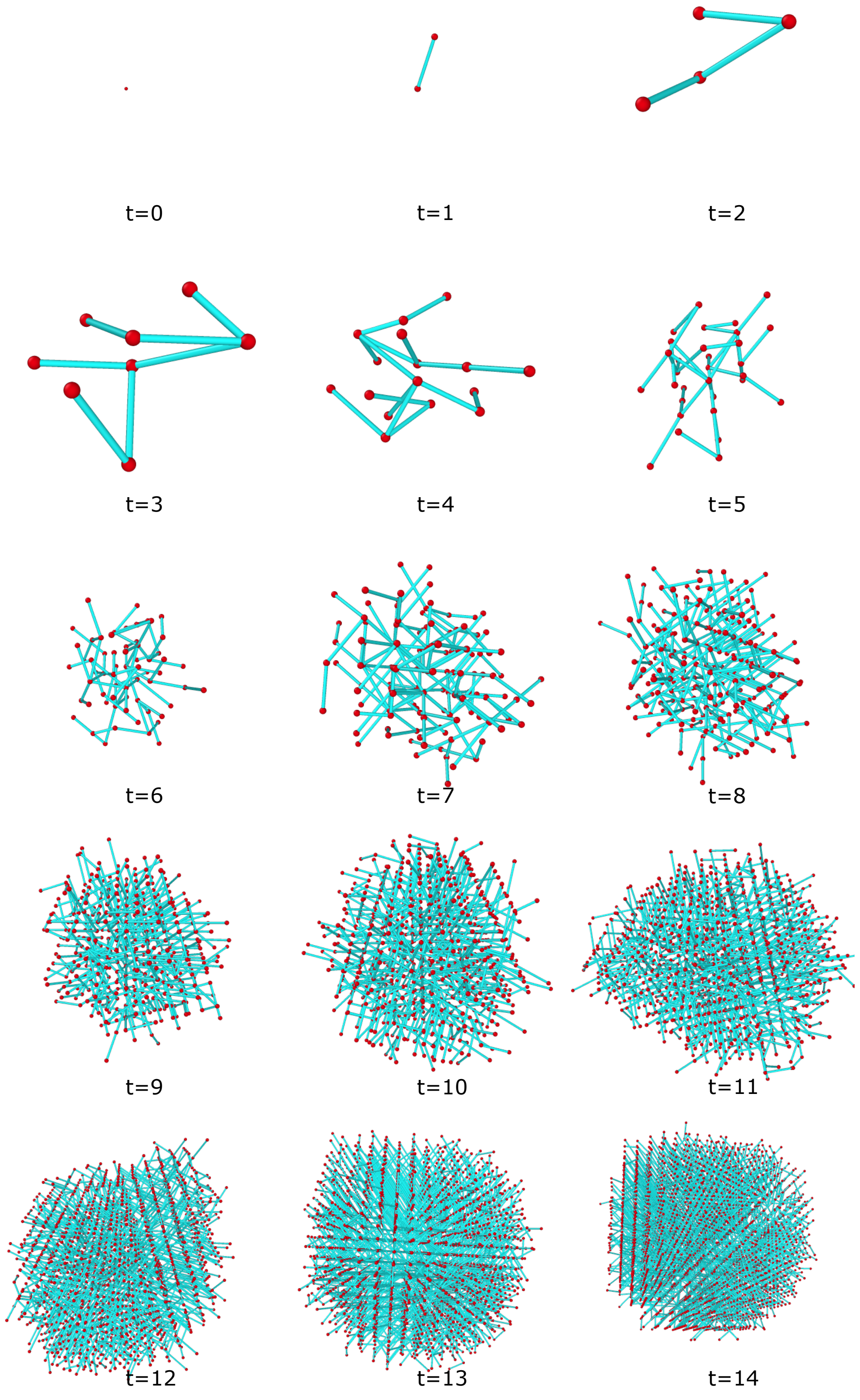


**Figure** S15: **Snapshots of the tumor evolution for** $\alpha = 1$. The red nodes are the cells and the blue edges denote the parent-child relationship.
